# Conformational diversity in poly-HAMP arrays and its implications for signal transduction

**DOI:** 10.64898/2026.02.18.704330

**Authors:** Murray Coles, Carolin P. Ewers, Reinhard Albrecht, Mikel Martinez-Goikoetxea, Malgorzata Orlowska, Jörg Martin, Andrei N. Lupas, Marcus D. Hartmann, Stanislaw Dunin-Horkawicz

**Affiliations:** Department of Protein Evolution, Max Planck Institute for Biology Tübingen, Max-Planck-Ring 5, 72076 Tübingen, Germany; Institute of Evolutionary Biology, Faculty of Biology, Biological and Chemical Research Centre, University of Warsaw, Żwirki i Wigury 101, 02-089 Warsaw, Poland; Interfaculty Institute of Biochemistry, University of Tübingen, Auf der Morgenstelle 34 72076 Tübingen, Germany

## Abstract

Prokaryotic transmembrane receptors are built around a helical coiled-coil backbone, with specialized sensory, modulatory, and effector domains arranged along its length. The modulatory HAMP domain forms a four-helix parallel coiled coil that is structurally integrated into this backbone, typically connecting transmembrane segments with downstream cytosolic domains. In many systems, HAMP domains have been shown to transduce signals through axial rotation of their helices; however, it is not clear how broadly applicable this mechanism is. Here, we describe two families of soluble chemoreceptors and sensory kinases that contain long arrays of concatenated HAMP domains, which we term poly-HAMP. Although their poly-HAMP arrays have clearly evolved independently, both families share many sequence features consistent with convergence on a similar functional system. We determined the crystal structures of 4-HAMP and 6-HAMP segments from the poly-HAMP array of histidine kinase HskS of *Myxococcus xanthus*, revealing unusually tight packing between adjacent domains and conformational patterns compatible with the rotational signaling model. To assess whether these features are general and to define the broader conformational landscape of poly-HAMP arrays, we computed AlphaFold2 models of over 200 chemoreceptor- and kinase-associated arrays. The models were consistent with the HskS structures, yet revealed distinct preferences in the two array types: HAMP domains of chemoreceptor arrays were consistently predicted to adopt stable conformations along their lengths, whereas those of kinase arrays were predicted to be biased toward less favorable conformations. By modeling HAMP domains from kinase arrays in isolation from the neighboring domains, we show that they adopt alternative, stable conformations that are related to the array-embedded forms through axial helix rotation. Taken together, our results suggest that, despite their independent origins and structural diversity, HAMP-containing systems ranging from the poly-HAMP arrays studied here through multi-HAMP architectures to canonical single-HAMP receptors may have converged on the same conserved mode of signal transduction that involves axial helix rotation.

## Introduction

Coiled coils (CCs) are structural protein motifs formed by two or more α-helices that wrap around each other in parallel or antiparallel orientation, forming regular bundles (Lupas et al., 2017; Woolfson, 2023). Initially identified in structural proteins such as keratins, where they provide mechanical stability, coiled coils were later found to serve many functions beyond structural support (Hartmann, 2017; Truebestein et al., 2016). This functional versatility is well illustrated by prokaryotic signaling proteins such as sensor histidine kinases and chemoreceptors, which are built around coiled-coil backbones with functional domains arranged along them (Diensthuber et al., 2013; Ferris, Coles, et al., 2014; Ferris, Zeth, et al., 2014). In these proteins, the coiled-coil backbone not only acts as a structural scaffold but also plays a central role in signal transduction, with some functional domains consisting entirely of coiled coils. Among these CC-based domains, one of the best-studied is the HAMP domain, which forms a parallel four-helix coiled-coil bundle and functions as a signal modulator and connector between domains of different types (Dunin-Horkawicz et al., 2010; Hulko et al., 2006).

At the sequence level, coiled coils, including HAMP domains, are characterized by a repeating pattern of seven residues (Lupas et al., 2005). The positions within each repeat are labeled *a* through *g*, with positions *a* and *d* typically occupied by hydrophobic residues facing the core of the bundle. These two core residues interact via knobs-into-holes packing, in which a residue from one helix (the knob) fits into a cavity formed by the residues of the opposing helices (the hole) (Crick, 1953). Unexpectedly, the first structure of a HAMP domain from the *Archaeoglobus fulgidus* receptor Af1503 revealed a conformation in which all helices were axially rotated relative to the canonical knobs-into-holes packing register (Hulko et al., 2006). In this rotated state, a third residue is incorporated into the hydrophobic core, resulting in a conformation known as x-da packing. While this extended packing is common in antiparallel coiled coils and is achieved by axial rotation of all helices in the same direction (Szczepaniak et al., 2014), in a parallel HAMP structure, it requires the input (N-terminal) and output (C-terminal) helices to rotate in opposite directions. This observation led to a signaling mechanism known as the “gearbox” model, in which HAMP interconverts between the knobs-into-holes and x-da states (Hulko et al., 2006). This “gearbox” model is consistent with findings from structural and functional studies in diverse HAMP-containing systems (Airola, Watts, Bilwes, et al., 2010; Ferris et al., 2011, 2012; Ferris, Zeth, et al., 2014; Klose et al., 2014; Mondéjar et al., 2012).

While most sensor kinases and chemoreceptors contain a single HAMP domain, typically connecting the membrane-anchoring region to downstream effector domains, some contain multiple copies. In these proteins, HAMP domains are either interspersed with other domains or arranged in poly-HAMP arrays consisting of many tightly connected HAMP units. Sequence analyses have shown that poly-HAMP arrays evolved through the repetition of di-HAMP modules, each composed of two types of HAMP domains: canonical ones, resembling the Af1503 HAMP, and non-canonical (or divergent) ones, which adopt a similar structure but share little sequence similarity with canonical domains (Airola, Watts, & Crane, 2010; Dunin-Horkawicz et al., 2010). Both canonical and non-canonical HAMP sequences exhibit features compatible with knobs-into-holes packing, though they differ in the nature of their alternative rotated states. We have previously proposed that while canonical HAMPs transition between knobs-into-holes and x-da states, non-canonical ones alternate between knobs-into-holes and da-x packing (Winski et al., 2024). Both the x-da and da-x states incorporate a third residue into the hydrophobic core but result from opposite rotations of the input and output helices. In canonical HAMP domains, the transition to x-da packing involves counterclockwise rotation of the input helices and clockwise rotation of the output helices. In contrast, in non-canonical HAMPs, da-x packing arises from the reverse: clockwise rotation of the input helices and counterclockwise rotation of the output helices. Therefore, placing the two HAMP types in tandem creates a self-contained toggle: the first reverses the sense of rotation, and the second reverses it again before passing the signal to the next di-HAMP module. This configuration allows the formation of extended arrays, which may otherwise stall if composed of only one HAMP type.

Multi-HAMP receptors constitute another class of proteins that contain more than a single HAMP domain. Unlike poly-HAMP arrays, which can comprise dozens of HAMP domains arranged in repeating, highly similar, and tightly connected di-HAMP modules, multi-HAMP receptors have fewer and more diverse HAMP domains. These domains are separated by linkers of variable length, ranging from short helices to entire domains such as PAS. A well-studied example of a multi-HAMP protein is the soluble receptor Aer2 (Airola, Watts, Bilwes, et al., 2010; Anaya et al., 2022), which contains five HAMP domains: three located at the N-terminus, followed by an oxygen-sensing PAS domain, and two additional HAMP domains. Another example is HtrII, a two-HAMP membrane protein that forms a light-sensing complex with sensory rhodopsin II (J. Wang et al., 2012).

Although the overall domain organization of poly-HAMP arrays is known, their biological roles and detailed structural properties remain largely uncharacterized. In bacteria, the HAMP-rich histidine kinase HskS from *M. xanthus* has been implicated in developmental regulation (Shi et al., 2008), while in fungi, deletions or mutations within poly-HAMP regions affect osmotic stress responses and fungicide resistance (El-Mowafy et al., 2013; Randhawa et al., 2016). This functional diversity prompted us to investigate whether the conformational plasticity of poly-HAMP arrays reflects a shared signaling mechanism. To address this question, we first performed sequence-based identification and classification of poly-HAMP arrays, followed by crystallographic characterization of the first six HAMP domains of the HskS poly-HAMP array. Given the high accuracy of AlphaFold2 in modeling coiled-coil domains (Madaj et al., 2025), including HAMP domains in different states (Winski et al., 2024), we then used it to predict the structures of more than 200 poly-HAMP arrays. By modeling these arrays both as entire assemblies and as individual HAMP domains separated from the array context, we not only generalized observations derived from experimental structures but also revealed alternative conformational states compatible with the “gearbox model”.

## Results and Discussion

### Identification and classification of poly-HAMP arrays

We identified poly-HAMP arrays by scanning the NCBI sequence database with profiles derived from canonical and non-canonical HAMP sequences from a manually curated set of representative arrays (see Methods). The resulting sequences were combined with HAMP domains from typical single-HAMP receptors and clustered based on pairwise sequence similarity (Figure 1A).

**Figure 1.**
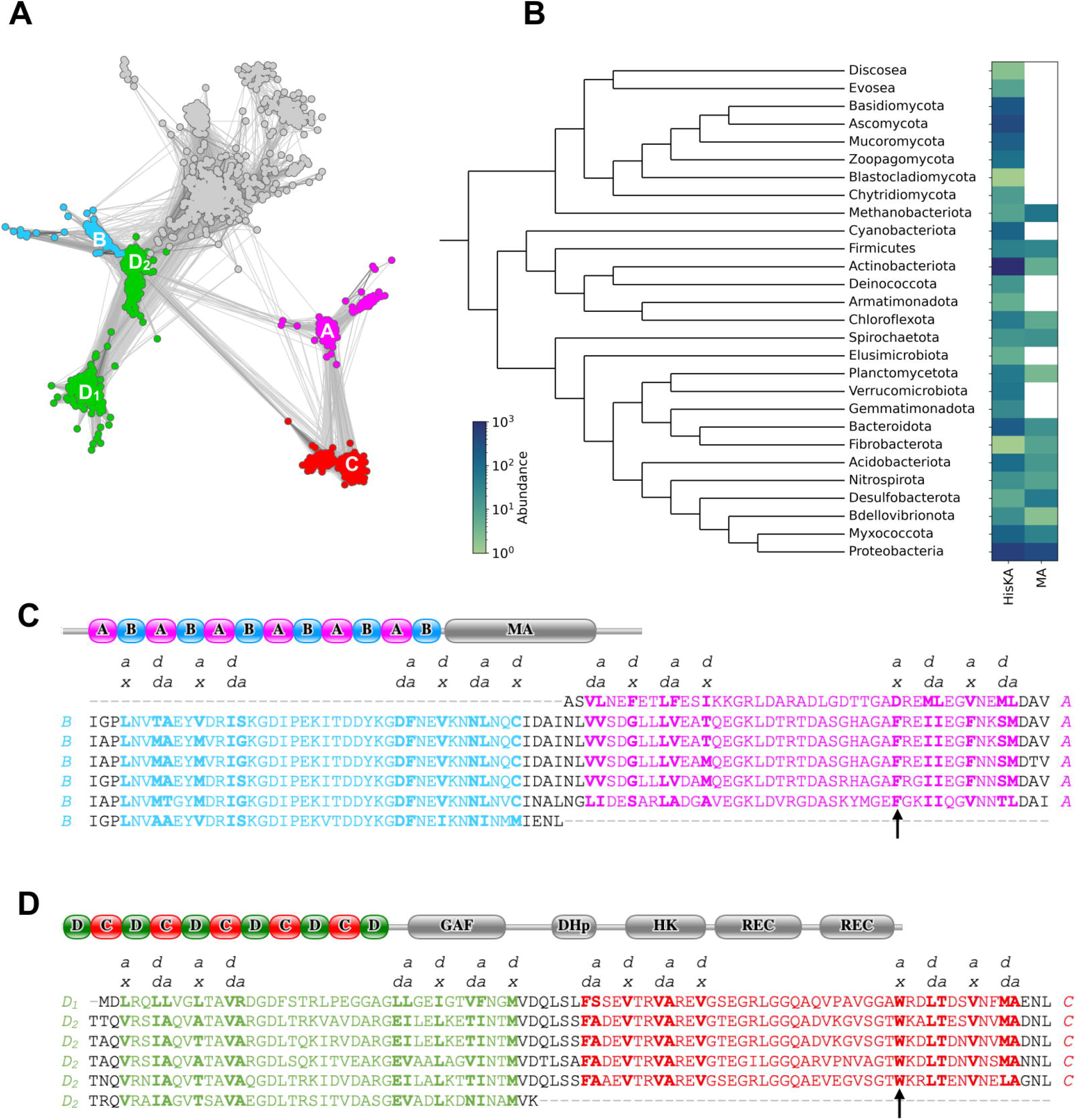
Sequence analysis of poly-HAMP array. (A) Cluster map of individual HAMP domain sequences, shown as circles positioned according to their pairwise BLAST similarity. Domains from poly-HAMP arrays are colored according to type (labeled A to D), while domains from single-HAMP receptors are shown in grey. (B) Phylogenetic distribution of phyla encoding either type of poly-HAMP arrays. (C) Alignment of di-HAMP modules from an exemplary chemotaxis-associated array (WP_274427672), with canonical and non-canonical HAMP domains colored in blue and purple, respectively. For each HAMP type, the knobs-into-holes and rotated (x-da or da-x) registers are indicated above. Residues contributing to the hydrophobic core are shown in bold. The downstream methyl-accepting (MA) domain is indicated. (D) Equivalent depiction for a kinase-associated array (AEW99355), with canonical and non-canonical HAMP domains shown in green and red, respectively. The sensory GAF domain, kinase domains (DHp and CA), and receiver (REC) domains are indicated.

The resulting cluster map revealed four HAMP groups, labeled A through D, that define two distinct types of poly-HAMP arrays. The first type, consisting of HAMPs A and B, is found upstream of the methyl-accepting domains characteristic of chemoreceptors. The second type, comprising HAMPs C and D, occurs upstream of multi-domain modules that include a GAF domain, histidine kinase domains (DHp and catalytic), and one or more receiver domains. The non-canonical HAMP domains (groups A and C) are distinct from each other and from all other sequences, whereas the canonical ones (groups B and D) are more similar to each other and to HAMPs from single-HAMP receptors, such as Af1503. Notably, canonical HAMPs in group D can be further subdivided into D1 and D2 subgroups, with D1 HAMPs consistently occupying the first position in the array.

Poly-HAMP arrays consist of repeating di-HAMP modules that are often nearly identical within a single array (Figure 1C and D), indicating an ongoing amplification process. Although chemotaxis- and kinase-associated arrays are composed of clearly distinct HAMP domain types, suggesting independent evolutionary origins, both share strikingly similar features. These include hydrophobicity patterns consistent with knobs-into-holes, x-da, and da-x conformations, identical inter-HAMP linker lengths, and conserved glycine residues at HAMP-HAMP junctions, which likely reflect convergence on a shared interdomain packing mode. Finally, the phylogenetic distribution of poly-HAMP arrays (Figure 1B) reveals their broad taxonomic spread, including possible horizontal gene transfer events between bacteria and eukaryotic lineages such as fungi and *Amoebozoa*.

### Experimental structure of kinase poly-HAMP

To directly characterize how HAMPs pack and what conformations they assume in poly-HAMP arrangements, we determined crystal structures of the first four and first six HAMP domains from the *M. xanthus* HskS kinase (MXAN_0712; UniProt: Q1DEE3), each fused to a C-terminal GCN4 adaptor to improve stability (Figure 2) (Deiss et al., 2014; Hernandez Alvarez et al., 2008). The full-length HskS kinase contains 21 HAMP domains, with the first belonging to the D1 subgroup of canonical HAMPs, followed by an alternating array of non-canonical (group C) and canonical (subgroup D2) domains. In the crystals of the 6-HAMP construct, we identified four chains in the asymmetric unit (ASU), belonging to three 6-HAMP dimers: two chains formed a dimer within the ASU, while the other two formed dimers through crystallographic symmetry. In contrast, the shorter 4-HAMP construct yielded a single dimer in the ASU. In total, we thus obtained three dimeric structures from the six-domain construct and one from the four-domain construct.

**Figure 2.**
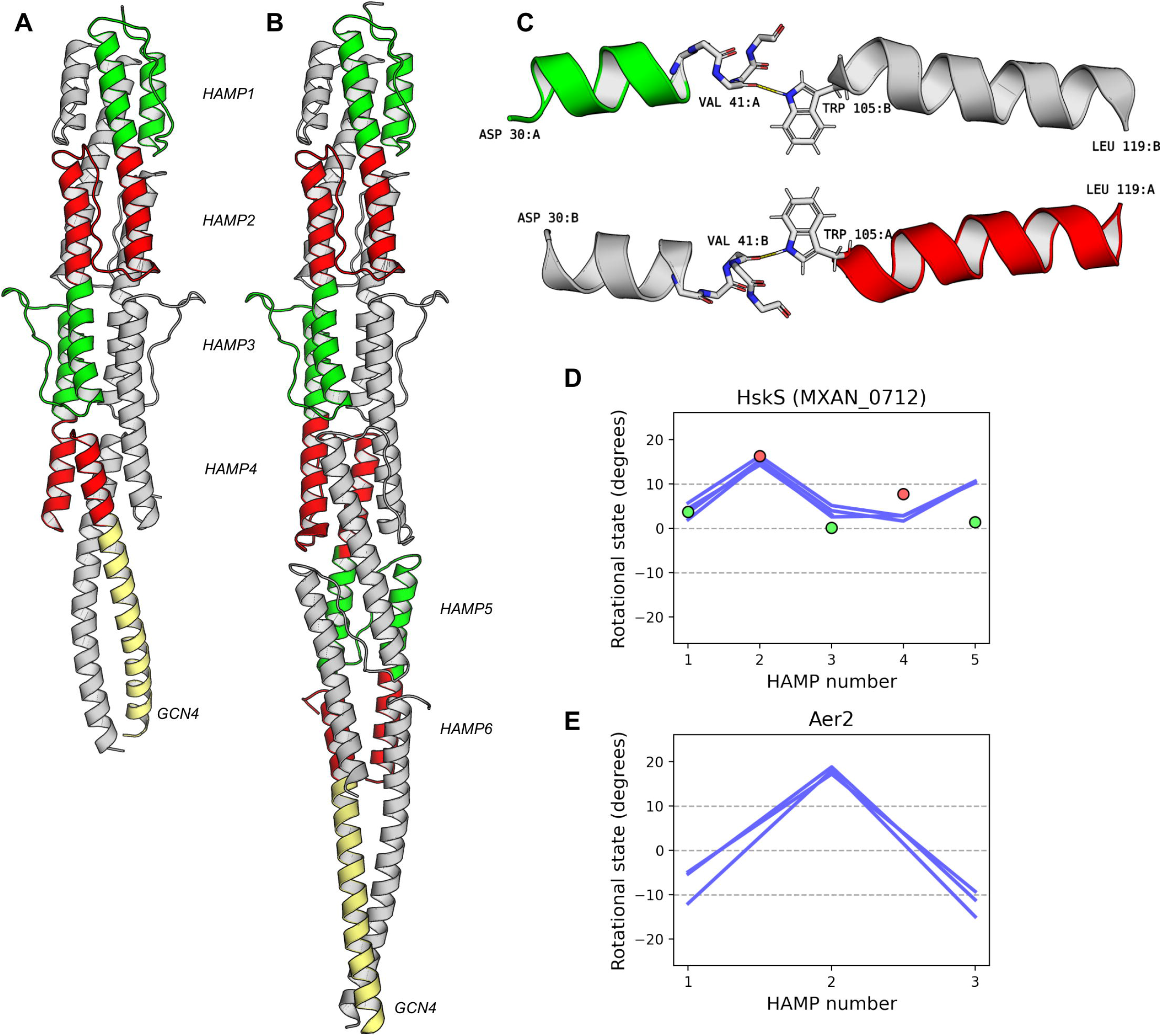
Experimental structures of poly-HAMP array fragments from the HskS hybrid kinase. Canonical and non-canonical HAMP domains are shown in green and red, respectively, with the C-terminal GCN4 stabilizing adaptor in yellow. For clarity, chain A is colored and chain B is shown in gray in panels A-C. (A) Structure of the four-HAMP construct. (B) Structure of the six-HAMP construct as observed in the asymmetric unit. (C) Close-up of the canonical-to-non-canonical interface showing hydrogen bonding between tryptophan 105 side chain and the backbone oxygen of valine 41. For clarity, only N-terminal and C-terminal helices of canonical and non-canonical HAMP, respectively, are shown. (D) Conformations of the first five and first three HAMP domains from the experimental structures (shown in blue), compared to the corresponding AlphaFold2 model (colored dots indicating domain type). Deformed terminal domains were omitted. (E) Conformations of the first three HAMP domains from the Aer2 multi-HAMP array.

Inspection of the four-domain construct revealed that the last HAMP domain deviated significantly from known HAMP structures, displaying an unstructured interhelical linker and distorted coiled-coil bundle. We suspected that this unusual conformation might be an artifact resulting from the fusion to the GCN4 adaptor. This suspicion was confirmed in the six-domain construct, where the first five HAMP domains, including the domain that previously appeared partially unstructured and misfolded, adopted their expected canonical conformations, while the final domain exhibited the same deviation as seen in the shorter construct. Given that the first (N-terminal) HAMP domains in both constructs are highly similar, we concluded that the influence of the GCN4 adaptor is primarily exerted on its adjacent domains (Hartmann et al., 2009). Consequently, for further analysis, we focused on the first three and five HAMPs from the four- and six-domain constructs, respectively.

Both constructs adopt a similar dimeric architecture in which all HAMP domains align along a common 2-fold symmetry axis. Each HAMP monomer consists of a pair of parallel helices connected by an unstructured segment. This interhelical linker is tightly packed into the interhelical groove through a combination of hydrophobic and polar interactions, and the monomers dimerize to form a four-helix coiled coil. The third HAMP in the array differs from the others in two respects. First, a proline residue in the N-terminal input helix induces a characteristic kink. Second, an insertion in the interhelical linker gives rise to a beta-hairpin structure at its C-terminal end. These two features are strongly conserved in the third HAMP of HskS-like poly-HAMPs, although their functional role remains unclear.

In the context of the array, individual domains are concatenated such that the output C-terminal helix of each HAMP is continuous with the input N-terminal helix of the next domain. Likewise, the N-terminal helix of each HAMP is nearly coaxial with the C-terminal helix of the preceding HAMP, giving the array the appearance of a continuous, supercoiled four-helix bundle. In single HAMP domains, each monomer consists of two helices of equal length, offset by one helical turn. As a result, the dimeric unit forms a central four-helix bundle flanked by short two-helix segments. In poly-HAMP arrays, however, these segments overlap completely, bringing neighboring domains into direct contact via backbone hydrogen bonds between the capping motifs of the respective helices. This close packing provides a rationale for the conservation of glycine residues in these motifs (Figure 1), as also observed at the HAMP2-HAMP3 junction in the Aer2 multi-HAMP module (Airola, Watts, Bilwes, et al., 2010).

The alternating arrangement of canonical and non-canonical HAMP domains gives rise to two distinct types of interdomain interface. In the canonical-to-non-canonical interface, a conserved tryptophan located at the N-terminal end of the C-terminal helix in the non-canonical HAMP forms a hydrogen bond with the backbone at the C-terminal end of the N-terminal helix in the upstream canonical HAMP (Figure 2C). The reverse, non-canonical-to-canonical interface is less specific, with a hydrophobic residue, most commonly isoleucine or valine, occupying the position of the tryptophan (Figure 1). The difference between these two interface types is also reflected in the geometry of the C-terminal capping motifs in the input helices. Whereas in non-canonical HAMPs, the N-terminal helix terminates in a classic six-residue Schellman motif (Aurora et al., 1998), in canonical HAMPs it adopts a less common seven-residue form (Datta et al., 1999). The resulting shift in local angles appears to facilitate interaction with the tryptophan in the downstream non-canonical HAMP.

The tight packing of HAMP domains observed in the experimental structures is reflected not only in the sequences of kinase-associated arrays but also in those of chemoreceptor-associated arrays (Figure 1), except that in the latter the conserved tryptophan in the canonical-to-non-canonical interface is replaced by a bulky hydrophobic residue, most commonly phenylalanine. This substitution reduces the ability of the interface to form the hydrogen bonds described above.

Finally, we analyzed the conformations of individual HAMP domains in the two experimental structures using Crick’s parametrization, which enables precise quantification of coiled-coil structures (Szczepaniak et al., 2021). For HAMP domains, the rotations of the input (N-terminal) and output (C-terminal) helices relative to the bundle axis were calculated separately, and the global rotation coefficient was defined as the difference between them (Winski et al., 2024). This coefficient places a given HAMP structure on a continuum of rotational states, with negative, zero, and positive values corresponding to x-da, knobs-into-holes, and da-x packing, respectively. This approach has the advantage of capturing the rotational state independently of the degree of bundle supercoiling, allowing fair comparisons across different HAMP types.

Application of this procedure revealed that the structures obtained from the four- and six-domain constructs are highly similar, with only minor differences between corresponding domains. The first and third canonical HAMPs in the array adopt essentially the same knobs-into-holes conformation, whereas the intervening non-canonical HAMP adopts a da-x packing state (blue lines in Figure 2D). This alternating pattern supports the “gearbox” model, according to which neighboring HAMP domains must operate in distinct conformational ranges. However, the downstream HAMPs adopt unexpected conformations that break this pattern. We suspect that this reflects a gradually increasing influence of the C-terminal GCN4 fusion along the array: the C-terminal HAMP is most affected, displaying an unstructured linker and distorted handedness, whereas upstream domains appear structurally intact but are shifted toward unusual rotational states, such that only the first three match the conformations expected within an array. Indeed, an AlphaFold2 model of the first five HAMPs without the GCN4 fusion extends this alternating arrangement to the downstream domains (dots in Figure 2D).

The structure of the HskS array fragment enabled us to describe the atomic details of HAMP-HAMP interactions and revealed evidence for an alternating pattern of conformations. It also expanded the set of canonical HAMP domains known to adopt the knobs-into-holes state (previously limited to examples such as CpxA (Mechaly et al., 2014), VicK (C. Wang et al., 2013), and the Af1503 A291F mutant (Ferris et al., 2011)), as well as the set of non-canonical domains known to adopt the da-x conformation which had so far been observed only in the second HAMP domain of the Aer2 multi-HAMP array structure (Figure 2E) (Airola, Watts, Bilwes, et al., 2010). However, this structure also raised new questions about whether these are general properties shared by kinase- and chemoreceptor-associated arrays and what conformational rearrangements occur during signal transduction. To tackle these questions, we modeled over 200 poly-HAMP arrays and their corresponding isolated HAMP domains with AlphaFold2 (Figure 3) and quantified their rotational states using the same procedure as for the crystallographic models (see Methods). Below, we show that modeling HAMPs both in isolation and in the context of their neighboring domains reveals alternative conformations of canonical and non-canonical HAMP domains that are compatible with the rotational “gearbox” model.

**Figure 3.**
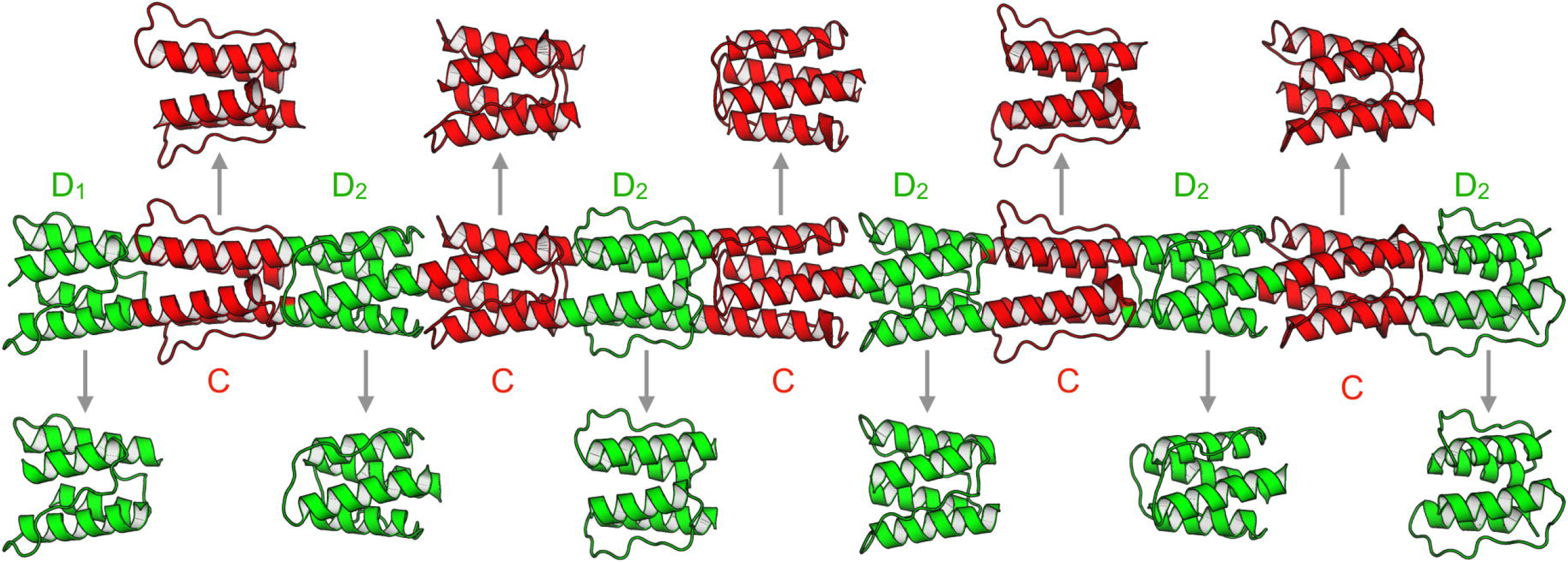
Predicting alternative HAMP conformations. The AlphaFold2 model of the exemplary kinase-associated poly-HAMP array (AEW99355) is shown in the center, flanked by models of isolated canonical and non-canonical HAMP domains below and above, respectively. In all structures, canonical and non-canonical domains are colored green and red, respectively. As described in the text, HAMP domains modeled in isolation and within arrays adopt distinct rotational states that are related by axial helix rotation, consistent with the gearbox model.

### Conformations of isolated HAMP domains

In previous work, we showed that AlphaFold2 models of HAMP domains adopt conformations closely matching those observed in corresponding experimental structures (Winski et al., 2024). Since most of these structures were crystallographic, we hypothesize that an isolated HAMP domain modeled in AlphaFold2 without neighboring domains may preferentially adopt a resting-like conformation. Based on this idea, we generated AlphaFold2 models of 11 individual HAMP domains isolated from an exemplary kinase-associated array (Figure 4A, top panel). These models revealed an alternating pattern of conformations, with canonical HAMPs consistently adopting the x-da state and non-canonical HAMPs adopting either the knobs-into-holes state or the da-x state. The same pattern was observed across isolated HAMP domains obtained from a diverse set of 120 kinase-associated arrays, where canonical HAMPs consistently adopted an x-da conformation, while non-canonical HAMPs sampled two alternative states (knobs-into-holes or da-x; Figure 4A, bottom panel). Notably, among non-canonical HAMPs with identical or nearly identical sequences, both the positively rotated da-x and the knobs-into-holes conformations were observed, suggesting that these reflect intrinsic conformational flexibility of individual domains rather than sequence-determined structural subtypes. A similar alternating pattern was observed in 99 chemotaxis arrays (Figure 4C), where canonical and non-canonical HAMPs adopt negatively and positively rotated states, respectively. However, these arrays lacked the distinct conformational alternatives observed in non-canonical HAMPs of kinase arrays.

**Figure 4.**
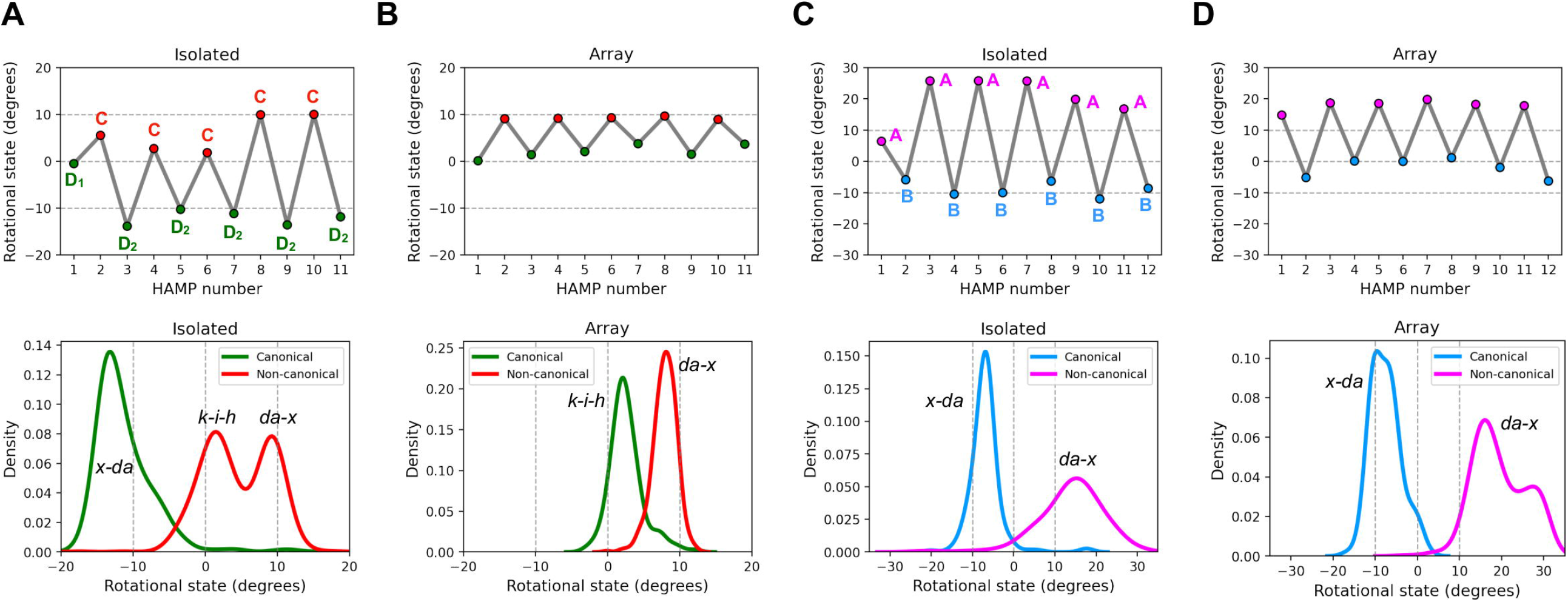
Comparison of isolated and arrayed HAMP conformations. (A) Rotational states of isolated HAMP domains from the kinase-associated array AEW99355 (top) and distribution of rotational states across isolated HAMP domains from 120 kinase-associated arrays (bottom). Rotational states corresponding to canonical knobs-into-holes packing (k-i-h) and rotated x-da and da-x packings are indicated. (B) Same representation as in (A), but for HAMP domains extracted from complete array models. Panels (C) and (D) show corresponding representations for the chemotaxis-associated array WP_274427672 and for a representative set of 99 other arrays of this type.

Modeling of individual HAMP domains highlighted an alternating zig-zag pattern of resting conformations in both kinase- and chemotaxis-associated poly-HAMP arrays (Figures 4A and 4C). Despite this shared organization, the conformational ranges sampled by canonical and non-canonical HAMPs differ between the two array types, with both x-da and da-x states shifted toward higher rotational values in chemotaxis arrays than in kinase arrays. This suggests that, like the characteristic hydrophobicity patterns and conserved HAMP–HAMP linker lengths, the alternating pattern represents a convergent solution that arose independently in kinase and chemotaxis arrays to satisfy the constraints of a common signaling mechanism.

### HAMP conformations in poly-HAMP arrays

According to the gearbox model, canonical and non-canonical HAMPs in poly-HAMP arrays should operate within two partially overlapping rotational ranges. Both share the knobs-into-holes state but differ in their alternative conformations, adopting x-da and da-x packing, respectively. In models of isolated non-canonical HAMPs from kinase-associated arrays, we observed the two expected conformations corresponding to the knobs-into-holes and da-x states (Figure 4A). In contrast, canonical HAMPs consistently adopted a single conformation matching the negatively rotated x-da state, as seen in single-HAMP receptors such as Af1503. We hypothesized that modeling these HAMPs in the context of their neighboring domains might better capture interdomain interactions and trigger conformations not observed in isolated models (Figure 3). Indeed, modeling 120 full-length kinase-associated arrays revealed a marked shift (Figure 4B): canonical HAMPs now adopted the knobs-into-holes conformation not seen in isolation, whereas non-canonical HAMPs that had previously remained in the knobs-into-holes state in isolation shifted into the positively rotated da-x state.

We also noted distinct behavior of D1-type HAMP domains, which occur exclusively at the beginning of kinase arrays (Figure 1). Whereas D2-type canonical and C-type non-canonical HAMPs adopted distinct conformations in their isolated and array-embedded forms, D1-type HAMPs consistently adopted the same rotated x-da conformation regardless of modeling context (Figure 5). We hypothesize that these N-terminal HAMP domains may act as a stabilizing cap that modulates the responsiveness of the downstream array.

**Figure 5.**
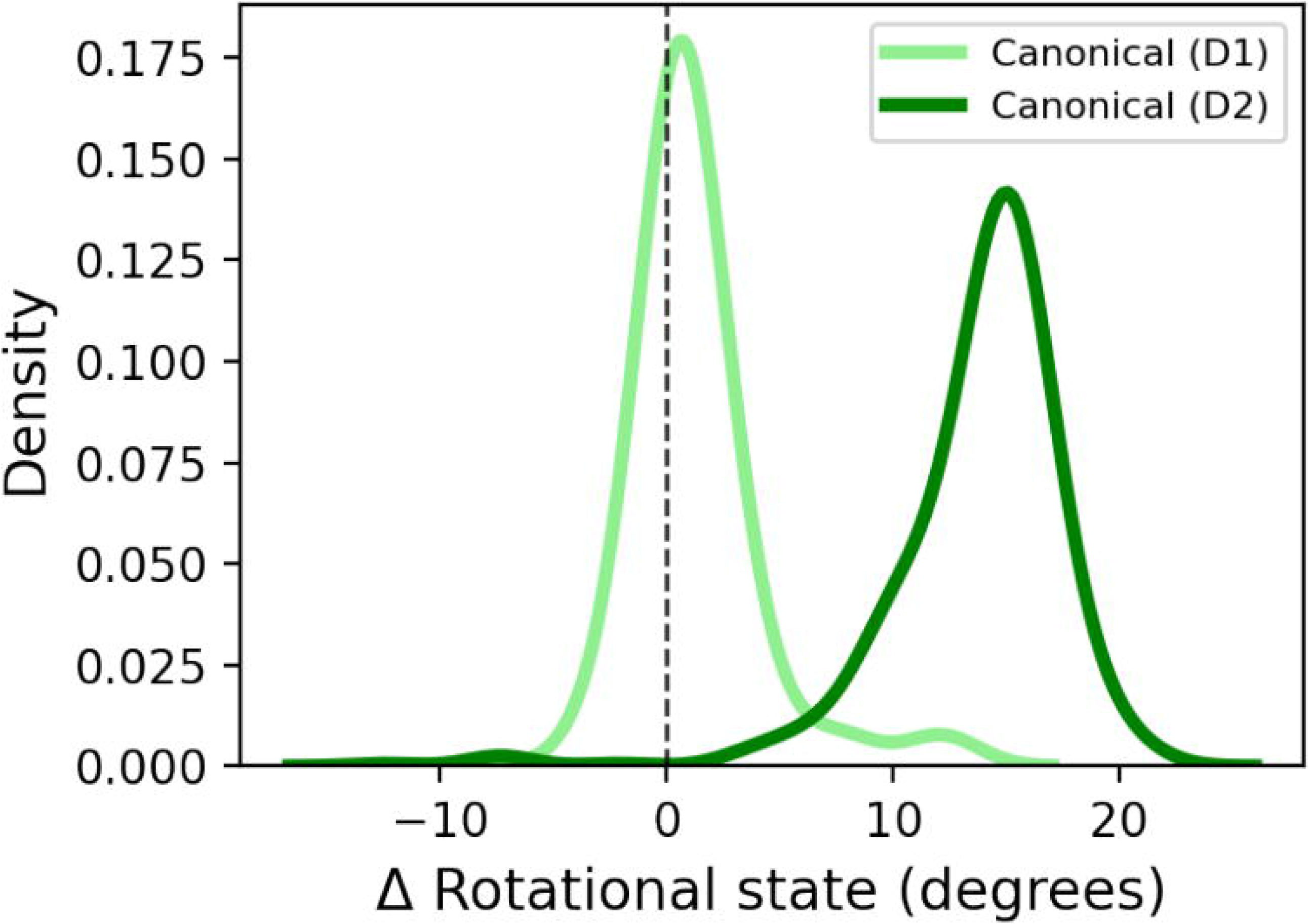
Conformational stability of the first and subsequent canonical HAMPs in kinase-associated arrays. The plots show the difference in rotational states between isolated and arrayed models for canonical D1 HAMPs (light green) and canonical D2 HAMPs (green). A value of zero indicates that the isolated and arrayed structures adopt the same conformation.

We employed the same procedure to explore possible alternative conformations in 99 chemotaxis-associated poly-HAMP arrays. However, in this case, we observed only modest differences between isolated and array-embedded models (compare panels C and D in Figure 4), rather than clearly distinct conformational states. Moreover, there was no clear difference between the N-terminal capping HAMPs that initiate the array and the downstream domains. This further emphasizes that, despite sharing a similar zig-zag pattern of conformations, kinase- and chemotaxis-associated arrays may differ in the details of their conformational dynamics and signal propagation.

### Implications for signal transduction mechanism

In typical single-HAMP receptors, the HAMP domain receives signals such as piston-like motions (Falke et al., 2001) or axial rotations (Moukhametzianov et al., 2006) from the membrane-spanning domain and converts them into a form interpretable by the downstream, usually effector, domain. In contrast, HAMP domains in poly-HAMP arrays are positioned downstream of other HAMP domains, and the entire array typically lacks an upstream input domain (Dunin-Horkawicz et al., 2010). However, a recent study has shown that the N-terminal, positively charged extension of the *M. xanthus* FrzCD multi-HAMP chemoreceptor can bind to DNA (Jazleena et al., 2025), suggesting that such arrays may be capable of coupling inputs at their extreme N-terminus to downstream signaling outputs. We identified similar positively charged extensions in several chemoreceptors analyzed in this study, indicating that DNA-binding propensity may be a feature shared by some chemotaxis-associated poly-HAMPs.

Chaining HAMPs into arrays gives rise to a zig-zag conformational pattern, as seen both in the HskS experimental structure and in modeled arrays (Figures 2 and 4). Despite differences such as variable intra-HAMP linker lengths and the absence of repetitive di-HAMP modules, a similar pattern is observed in the structure of the first three HAMP domains of the multi-HAMP array of the Aer2 soluble chemoreceptor (Airola, Watts, Bilwes, et al., 2010) (Figure 2E). The first and third HAMPs adopt negatively rotated states matching the x-da packing expected for canonical HAMPs, whereas the second adopts the da-x packing typical of non-canonical ones. Interestingly, the sequence of the second Aer2 HAMP domain falls within group A HAMPs (Figure 1), which includes non-canonical domains of chemoreceptor arrays. This suggests that Aer2-like multi-HAMP arrays and the chemoreceptor poly-HAMP arrays analyzed here may, at least in part, share a common evolutionary origin. An alternating conformational pattern was also proposed for the HtrII two-HAMP photoreceptor (J. Wang et al., 2012). Although no experimental structures are available in this case, fluorescein probe accessibility scanning indicated opposite motions in the first and second HAMP domains, consistent with a zig-zag conformational pattern.

Despite general structural similarities suggesting a shared mechanism of signal transduction, kinase- and chemoreceptor-associated poly-HAMP arrays exhibit distinct behaviors. In the kinase family, pronounced conformational differences are observed between isolated and array-embedded forms, whereas in the chemoreceptor family, the two forms are very similar (Figure 4). We hypothesized that the conformational discrepancy in the kinase family may arise from conformational constraints imposed on HAMP domains embedded within an array. These constraints could stem from specific interdomain interactions unique to kinase-associated arrays, such as the conserved tryptophan-mediated hydrogen bonds (Figure 2C). In isolated models, where these contacts are absent, HAMP domains may relax into alternative, unconstrained conformations that are otherwise inaccessible within the array.

To test this, we analyzed AlphaFold2 pLDDT scores and Rosetta scores for isolated and arrayed domains. While the Rosetta score is often used as a proxy for relative stability (Alford et al., 2017), AlphaFold2’s pLDDT score primarily reflects model confidence, but it has also been associated with flexibility (Guo et al., 2022; Ma et al., 2023; Vander Meersche et al., 2025). The results obtained reveal a clear distinction between kinase and chemoreceptor families: in kinase arrays, isolated models exhibit consistently higher pLDDT scores and lower per-residue Rosetta scores, whereas in chemotaxis arrays, the differences are minimal (Figure 6). This suggests that HAMP domains in kinase arrays may populate more dynamic conformations, consistent with the dynamic bundle model, in which the HAMP domain transduces signals by switching between stable and dynamic states (Stewart, 2014; Sukomon et al., 2017; Zhou et al., 2011). A similar mechanism has been proposed for the Aer2 multi-HAMP receptor, where molecular threading analysis indicated increased conformational freedom of the second, non-canonical HAMP domain (Airola, Watts, Bilwes, et al., 2010).

**Figure 6.**
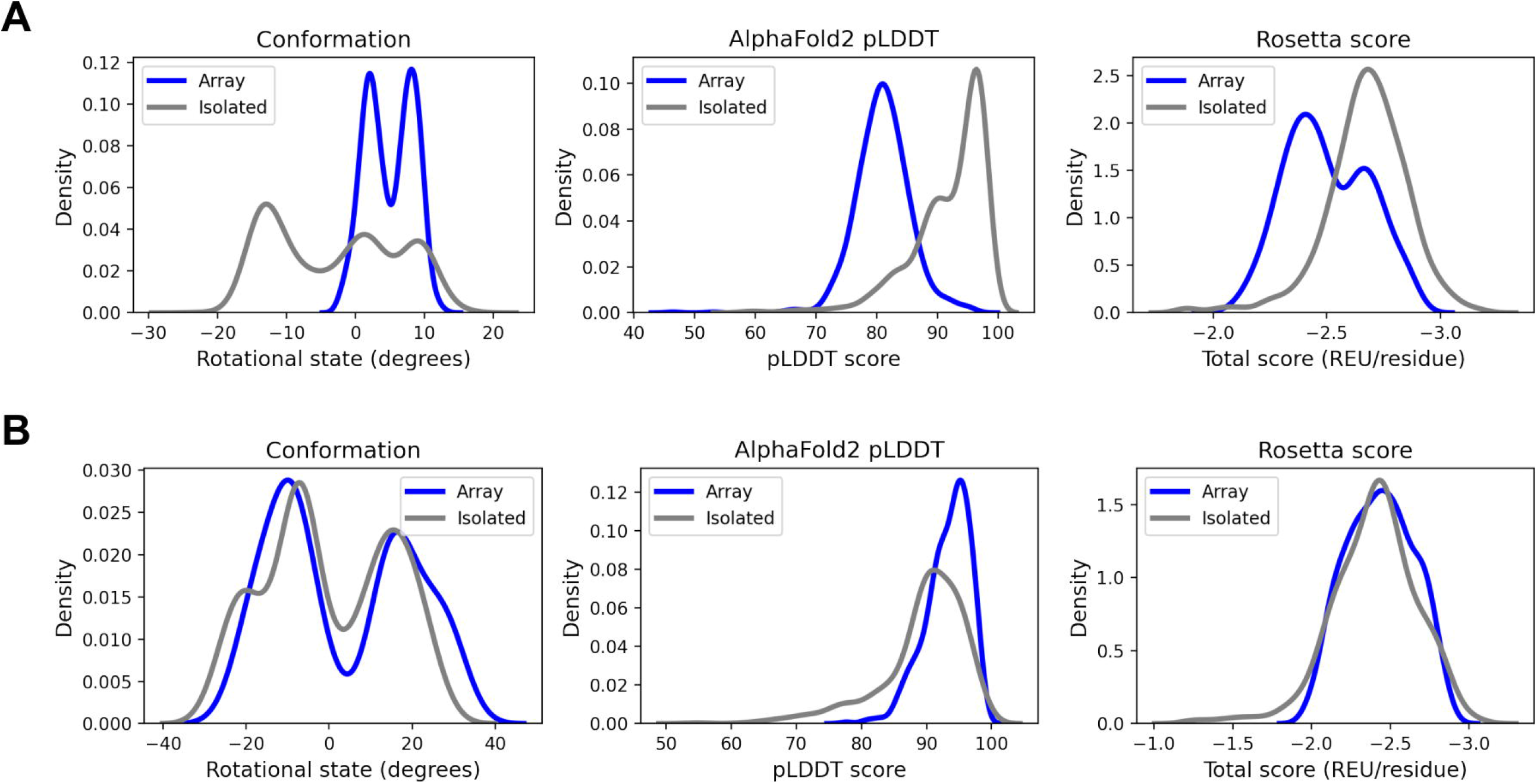
Comparison of AlphaFold2 pLDDT scores and per-residue Rosetta scores between arrayed and isolated HAMP models. (A) Distributions of conformational states, pLDDT scores, and Rosetta scores for arrayed (blue) and isolated (grey) HAMP domains from kinase-associated arrays. (B) Same analysis as in (A), shown for chemoreceptor-associated arrays.

Finally, we used molecular dynamics simulations (MD) of a canonical HAMP domain from a kinase-associated array to investigate the structural basis of conformational switching. The first simulation was initiated from a structure extracted from the AlphaFold2 model of the full array. This structure adopted an unrotated knobs-into-holes packing that, upon release from the constraints imposed by neighboring domains, rapidly transitioned to a negatively rotated x-da conformation (Figure 7). This relaxation involved not only rotation of the helices toward the x-da state but also a decrease in RMSD relative to the AlphaFold2 model of the isolated domain. In contrast, a simulation initiated from the isolated model of the same HAMP remained in the x-da state throughout. These results suggest that the discrete conformations obtained with AlphaFold2 can be connected through a continuous transition in MD, indicating that interconversion between these states is physically plausible once array constraints are released. While this analysis was limited to a single domain, it illustrates the potential of molecular dynamics to probe transitions between HAMP conformational states and motivates further investigation in this direction.

**Figure 7.**
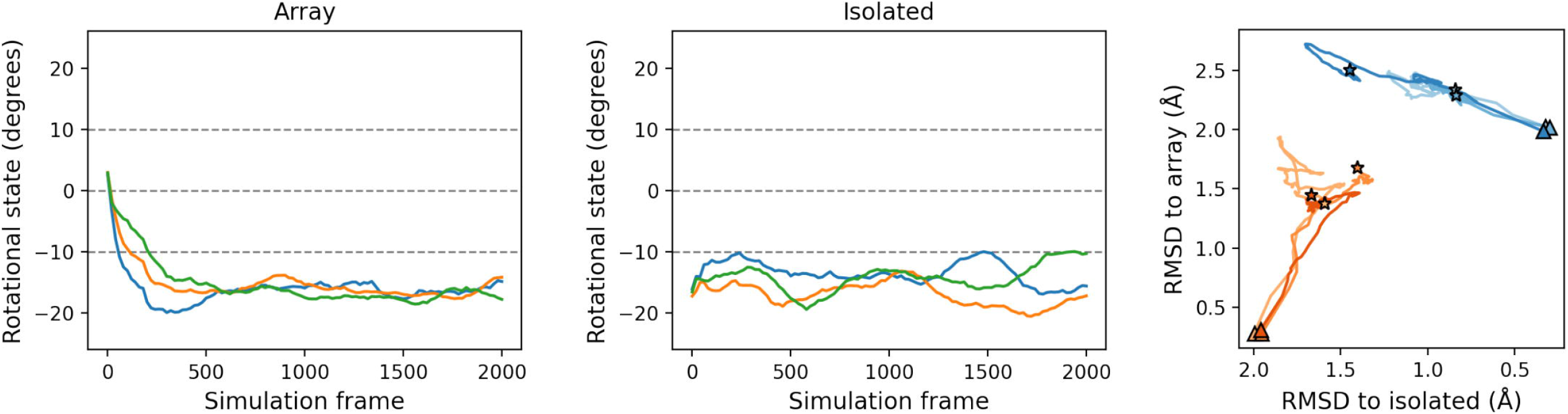
Molecular dynamics simulations of canonical isolated and arrayed HAMP domains. All simulations were performed in three independent replicas. The left and middle panels show the evolution of helix rotation angles over time for the arrayed and isolated forms, respectively. The right panel shows the RMSD trajectories from all replicas (note the reversed x-axis), with isolated and arrayed forms shown in shades of blue and orange, respectively.

## Conclusions

Based on the experimental structure of a poly-HAMP array and computational analyses, we propose that poly-HAMP arrays operate along a conformational continuum ranging from x-da to knobs-into-holes to da-x packing, with the first and last conformations predominantly accessible to canonical and non-canonical HAMPs, respectively. Given the length of these arrays, simultaneous conformational switching of all HAMP domains appears unlikely; we therefore propose that signal propagation may occur as waves of local relaxation to an alternative state. However, the nature of this propagation may be family-specific – in contrast to kinase arrays that may reside in a strained, more responsive state, chemoreceptor arrays adopt more relaxed conformations (Figure 6) and thus may require stronger stimuli to induce local switching.

While related multi-HAMP arrays, such as those of FrzCD, Aer2, and HtrII receptors, receive input signals at or near their N-terminal regions, poly-HAMP arrays appear to lack upstream sensory domains. Therefore, the role of signaling via poly-HAMP arrays remains unclear, raising the possibility that they act as tunable modulators through evolutionary changes in array length, for example via amplification of di-HAMP modules, or that individual HAMP domains may possess intrinsic ligand-recognition capabilities.

## Methods

### Sequence analysis

In 34 chemoreceptor and kinase sequences (Supplementary Table 1), HAMP domains were identified using HH-suite (Steinegger et al., 2019), pLM-BLAST (Kaminski et al., 2023), and Foldseek (van Kempen et al., 2024). Identified HAMP sequences were grouped by similarity using CLANS (Frickey et al., 2004), yielding nine clusters. Sequences in each group were aligned with PROMALS3D (Pei et al., 2008), and the resulting multiple sequence alignments were manually curated, including assignment of the coiled-coil register to the alignment columns. The curated alignments were used to build HMM profiles with HMMER 3.1b2 (Eddy, 2011). These profiles were then used to search the NCBI non-redundant database clustered at 90% sequence identity (NR90; version dated May 3, 2023) using hmmsearch with default parameters. The resulting matches were concatenated, yielding 5,501 proteins (4,778 histidine kinases and 723 chemoreceptors) containing poly-HAMP arrays with five or more consecutive HAMP domains. For visualization of their abundance and distribution (Figure 1B), a tree was constructed using GTDB taxonomy (Parks et al., 2020) for Bacteria and Archaea, and published classifications for Fungi (Spatafora et al., 2016) and other eukaryotes (Burki et al., 2020).

### AlphaFold2 modeling

In total, 120 kinase- and 99 chemoreceptor-associated poly-HAMP arrays were selected for modeling with AlphaFold2 (Supplementary Table 2) both as full-length arrays and as individual HAMP domains. Selection was based on sequence clustering of all identified HAMP domains using MMseqs2 (min_identity = 0.45, coverage = 0.5, cov_mode = 0, cluster_mode = 0) to obtain a diverse set; the corresponding full-length arrays containing these representative HAMPs were then retrieved. Modeling was performed with AlphaFold2 (version 2.3.1) (Jumper et al., 2021) in AlphaFold-Multimer mode (dimers), 5 predictions per AF2 model (a total of 25 predictions), and without templates. The models were sorted by ranking confidence (0.8\**ipTM + 0*.*2\**pTM), and the top-scoring model was chosen for subsequent analyses. To compare modeling confidence between HAMP domains modelled in isolation and in arrays, per-residue pLDDT scores for individual HAMP domains modelled in isolation and within arrays were extracted, averaged, and plotted as distributions (Figure 6, middle panel). These analyses were complemented by Rosetta 3.12 scoring (Leaver-Fay et al., 2011), where models were subjected to fast relax with coordinate restraints and evaluated using averaged per-residue Rosetta total energy scores computed with the ref2015 score function (Alford et al., 2017) (Figure 6, right panel).

### Coiled-coil parameters measurement

For quantifying conformations of isolated and arrayed HAMP domain models, we used Crick’s equations as implemented in the SamCC Turbo tool (Szczepaniak et al., 2021). A key requirement of this analysis is a consistent coiled-coil register assignment, which ensures that functionally corresponding regions are compared across analysed HAMP domains regardless of their type. As described in the “Sequence analysis” section, each query MSA used to identify HAMP domains in the NR90 database was manually assigned a coiled-coil register. These register annotations were propagated to all identified HAMP domains and used as a reference for measuring axial helix rotation of the input (N-terminal) and output (C-terminal) helices, as described in detail in our previous work (Winski et al., 2024).

### Cloning, expression and purification

DNA fragments encoding HskS’s N-terminus together with the first four (residues 1-212, “4-HAMP”) or the first six (residues 1-304, “6-HAMP”) HAMP domains were obtained from *M. xanthus* DK1622 genomic DNA (Kaiser, 1979) by PCR, using primers adding an NdeI restriction site to the start and an extension comprising an Eco31I site, a sequence encoding a Gly-Ser-Gly linker and an XhoI restriction site to the end. The PCR products were ligated to the NdeI and XhoI sites of pET28b(+), thereby adding an N-terminal His_6_ tag and a thrombin cleavage site. After treatment with XbaI and Eco31I, the excised fragments were cloned into pIBA-GCN4di, a vector derived from pASK-IBA2 (IBA Lifesciences GmbH) by N- and C-terminal insertions of a codon-optimized dimeric leucine zipper (GCN4 from *Saccharomyces cerevisiae*) while preserving its multiple cloning site (analogous to pIBA-GCN4tri (Hernandez Alvarez et al., 2008)). In the final constructs, the N-terminal His_6_-Tag was retained and the C-terminal 29 residue modified GCN4 sequence (MKQLEWKVEELLSKNYHLENEVARLKKLV) formed a continuous coiled coil in a register compatible with that of the last HAMP helix to which it was fused, including the GSG linker residues.

*Escherichia coli* C43 (DE3) cells (4-HAMP) or Rosetta2 cells (6-HAMP) were transformed with the constructs and cells were grown in LB medium supplemented with ampicillin. Protein expression was induced at an OD_600_ of 0.5 to 0.6 with anhydrotetracycline (0.25 μg/ml culture) at 25 to 28 °C for 6 to 8 h. Cells were harvested by centrifugation. We found that the presence of urea was advantageous for protein solubility and stability (Dines et al., 2007). Thus, the cell pellets were resuspended in lysis buffer (50 mM Tris, 300 mM NaCl, 1 mM MgCl_2_, 2 M urea, 0.5 mM dithiothreitol (DTT), 10% glycerol, pH 8, supplemented with DNaseI, lysozyme and phenylmethyl sulfonylfluoride). Lysis was performed with a French Press. Cleared lysate was loaded onto an equilibrated 10 ml Ni-NTA column. The column was washed with 10% elution buffer (as lysis buffer but containing 300 mM imidazole) and the protein was eluted with 100% elution buffer. Eluted protein was dialyzed against 20 mM Tris, 100 mM NaCl, 1.5 M urea, 1 mM MgCl_2_, 2 mM DTT, 5% glycerol, 10 mM imidazole, pH 7.8. White precipitate formed during dialysis, but the supernatant still contained sufficient amounts of the proteins which could be concentrated in 10 kDa spin concentrators. The purity of the isolated proteins was confirmed by SDS-PAGE.

### Crystallization, Data collection, and Structure Solution

Crystallization was performed at 20 °C via the vapor diffusion method. Initial trials were set up with a Honeybee 961 robot (Genomic Solutions Ltd.) by mixing 300-400 nl of protein solution and the same amount of reservoir solution on 96-well sitting-drop microplates, using commercially available screens (“Nextal”, Qiagen). Initial hit conditions were reproduced and refined in hanging-drop setups using EasyXtal plates (Qiagen). Best-diffracting crystals of 4-HAMP were obtained with a protein concentration of 5.3 mg/ml in condition G11 of the Classic II Screen (25% PEG 3350, 0.1 M BisTris pH 6.5, 0.2 M MgCl_2_). For phasing, heavy-atom derivatives of 4-HAMP crystals were obtained by soaking with platinum salts. Prior to flash-cooling, all 4-HAMP crystals were cryoprotected with 15% glycerol. Best-diffracting crystals of 6-HAMP were found at a protein concentration of 17 mg/ml in condition H12 of the Classic I Screen (12% PEG 20000, 0.1 M MES pH 6.5). These crystals were sensitive to aging and had to be harvested immediately after their formation for good diffraction. Cryoprotection was done with 20% PEG 300 and 0.4 M urea.

All data for 4-HAMP and 6-HAMP crystals were collected on beamline X10SA at the Swiss Light Source at 100 K. Derivative data were collected on a MarCCD 225-mm charge-coupled device (CCD) detector at the Pt L-III edge (peak), all native data were collected using a Pilatus 6M hybrid pixel detector (DECTRIS) at a wavelength of 1 Å. XDS was used for processing and scaling of the diffraction data, with the statistics given in Supplementary Table 3. Phasing of 4-HAMP crystals was done in two steps. First, initial phases were obtained following the Multiple Isomorphous Replacement with Anomalous Scattering (MIRAS) protocol with one native and two diammino platinum dinitrite derivative datasets. SHELXD (Sheldrick, 2008) was employed for heavy atom location as implemented in autoSHARP (Vonrhein et al., 2007), followed by substructure refinement using SHARP (de La Fortelle et al., 1997). After density modification with Solomon (Abrahams et al., 1996), the resulting map was still too noisy for manual or automated interpretation. For this reason, in a second step, we employed phased molecular replacement aided by the Solomon-improved phases as implemented in MOLREP (Vagin et al., 2010) with different HAMP domain models as search models. The first successful phased search was conducted with a two-HAMP module derived from Aer2 (Airola, Watts, Bilwes, et al., 2010), followed by a successful phased search employing a single AF1503 HAMP domain (Ferris et al., 2011) as a search model, thereby locating the first three HAMP domains of the construct. After initial manual model building in Coot (Emsley et al., 2004) and refinement with REFMAC5 (Murshudov et al., 1999), a further molecular replacement search with MOLREP could locate the C-terminal GCN4 adaptor using PDB entry 1GK6 (Strelkov et al., 2002) as a search model. The final structure comprises two chains in the ASU forming a single 4-HAMP dimer. Phasing of 6-HAMP crystals was performed by molecular replacement in MOLREP (Vagin et al., 2010) using the 4-HAMP structure as a search model, locating four chains in the ASU; these belong to three 6-HAMP dimers, two of which are built by 2-fold crystallographic symmetry. All structures were completed by cyclic manual modeling with Coot (Emsley et al., 2004) and refinement with REFMAC5 (Murshudov et al., 1999) with local NCS restraints. In both structures, apart from the N-terminal 25-30 residues, the linker in the last HAMP domains, and the C-terminal portions of the GCN4 adaptors, the full chain of all poly-HAMP molecules could be traced in the electron density. The refinement statistics are given in Supplementary Table 3; coordinates and structure factors were deposited in the PDB under accession codes 9TRJ (4-HAMP) and 9TRK (6-HAMP).

### Molecular dynamics simulations

The simulations were performed in GROMACS 2024.3 (Abraham et al., 2015; Van Der Spoel et al., 2005) using the CHARMM36m force field (Huang et al., 2017). Bond vibrations were constrained with the LINCS algorithm (Hess, 2008). Long-range electrostatics were treated with the PME method, and nonbonded interactions were computed using the Verlet cutoff scheme with a 1.2-nm real-space cutoff. A 2-fs integration time step was used. Systems were solvated with TIP3P water in a dodecahedral periodic simulation box with a minimum solute–box distance of 1.4 nm, and the net charge was neutralized with Na^+^/Cl^−^ ions as needed. Energy minimization was performed using the steepest-descent algorithm until the maximum force was below 300 kJ mol^−1^ nm^−1^. Following 1 ns equilibration in both the NVT and NPT ensembles, production simulations were run for 500 ns in triplicate. Temperature was controlled with the V-rescale thermostat, and pressure was maintained at 1 bar using isotropic coupling with the C-rescale barostat.

This initial set of simulations was carried out at 300 K using the arrayed and isolated AlphaFold2 models of the fifth HAMP of the CAA9318664 array. During the simulations, all systems remained globally stable in terms of RMSD, but simulations started from the arrayed model showed rapid transition to the rotational state seen in the isolated model. To slow this convergence, a second set of simulations was performed at 250 K for 200 ns in triplicate. Even at reduced temperature, the arrayed HAMP rapidly adopted the x-da state corresponding to the conformation observed in the isolated form (Figure 7).

## Acknowledgments

This work was supported by institutional funds from the Max Planck Society. S.D-H. was additionally supported by the National Science Centre, Poland (grant no. 2020/37/B/NZ2/03268). We thank Kerstin Bär for assistance with protein crystallization and the staff of beamline X10SA of the Swiss Light Source (PSI, Villigen, Switzerland) for technical support.

## Notes

### Competing Interest Statement

The authors have declared no competing interest.

