## Supplementary Table for "Conformational diversity in poly-HAMP arrays and its implications for signal transduction"

**Supplementary Table 1.** Poly-HAMP chemoreceptors and kinases whose HAMP domains were used as seed queries for searching the NCBI non-redundant sequence database.

| Kinases | A0A1N6G1D1, A0A4R4P399, A0A248K2C0, I4YVW6, A0A542ZIX7, A0A1U7JAJ8, A0A4Z0BGY9, A0A1G9HBX6, A0A3E0HHV0, A0A1N7CPU7, A0A3A9YLK8, K9PBI3, G0G0R7, D8FVA6, A0A0X3XHH5, A0A1H5PQZ5, A0A2W2M3F7, A0A2U1VYS9, B0UB08, A0A2D3U389, A0A1Q9RWB5, A0A0X3RTC7, A0A2A3HZJ6 |
| --- | --- |
| Chemoreceptors | Q2FN62, Q2FQJ9, Q2FQT7, Q74DM2, B9M9P4, Q3BU77, Q3BU80, A0A1S8RTT3, Q2FRC3, Q5H2U8, B2FI15 |

**Supplementary Table 2.** Poly-HAMP chemoreceptor and kinase arrays modeled in AlphaFold2 either as full-length arrays or as individual HAMP domains.

| Kinases | ABF13477, ABI63596, ACK51402, AEW99355, AKF79476, AOR06392, AOW78957, APR78731, APV48351, BAU86971, CAA9213639, CAA9276420, CAA9318664, CAA9514814, CAA9541585, CAD6583708, CAF9929585, CAG8476750, CAG8501670, CAH2352000, CAI5760408, CAJ65078, CDH51117, CDH57830, CDS55145, CUI03596, EFL09696, GDY41841, GET40449, HAK93061, HBE57325, HJE60676, KAA0680100, KAF2719822, KAF7332700, KAF7747413, KAF8308501, KAF8737078, KAG0199653, KAG8912441, KAG8920499, KAH8922829, KAH8924750, KAI0338803, KAI9749930, KAJ1945458, KAJ1957453, KAJ2455678, KAJ3014897, KAJ7025691, KAJ7714292, KFN46324, KJA22434, MBA2310897, MBA2348676, MBA2404959, MBA2794146, MBA3245812, MBA3466498, MBA3561234, MBA3594958, MBA3664223, MBA3684214, MBA4156052, MBC7174319, MBC7448616, MBC7893908, MBE3086571, MBF6568894, MBI1912230, MBI2378142, MBI2386612, MBI3313561, MBI3359409, MBI3558156, MBI4205782, MBI4348258, MBI4811967, MBI5438214, MBI5607008, MBK9001788, MBK9201114, MBN9394845, MBT3386106, MBT9495849, MBV52702, MBV8298408, MBV9866075, MBW3637220, MBX3187116, MBX3198964, MBX6388644, MCC7539601, MCG8634810, MCH8347281, MCJ7470134, MCU0534516, MPY94410, NRQ38315, NUO48611, NUO51383, OLD49681, OLE64068, QFG26374, RKP25275, TFW05571, TMA17525, TMA20250, TWV11213, WP_013130450, WP_029722888, WP_051794678, WP_092264814, WP_136535146, WP_141958392, WP_147304648, WP_228057440, XP_003889300, XP_036637199, XP_047801733 |
| --- | --- |
| Chemoreceptors | CUU50852, ESS73188, GAK55060, HBC93786, MBF0361290, MBK7877023, MBK8338217, MBM2814453, MBN2695045, MBN2695612, MBN2833792, MBS4050147, MCB1190930, MCB9496075, MCD6207395, MCG2797269, MCH2340873, MCM2335051, MCP5495817, MCP5502499, MCT4632982, NLL09390, NLO19448, NOT67721, NOU23061, NOX88659, NRB67684, NUN71418, NVM22399, OPZ36790, OQA77434, PKL59521, PKL60757, PZM89608, TMN88776, TQD27560, WP_008429341, WP_012631780, WP_013484833, WP_019897594, WP_020565711, WP_041362574, WP_071392859, WP_091471401, WP_109941520, WP_109966919, WP_110186788, WP_124551638, WP_128694700, WP_135279084, WP_140003491, WP_148602150, WP_155468411, WP_157202402, WP_159355165, WP_162299629, WP_166404993, WP_190259213, WP_202905785, WP_207688227, WP_207688826, WP_209620254, WP_210996399, WP_211531235, WP_214421110, WP_221063350, WP_221064443, WP_226636625, WP_228519770, WP_229716082, WP_230838595, WP_235811085, WP_236210111, WP_238916629, WP_239441545, WP_248342755, WP_248343955, WP_248345650, WP_248345651, WP_248345920, WP_254458424, WP_255332674, WP_256601726, WP_256613233, WP_257743022, WP_257743469, WP_258346527, WP_260610421, WP_260788041, WP_263393292, WP_263634519, WP_269745533, WP_274396358, WP_274427672, WP_275355203, WP_275356878, WP_275357030, WP_275357036, WP_277275308 |

**Supplementary Table 3.** Crystallographic Data Collection and Refinement Statistics

| Structure | **4HAMP** | **6HAMP** |
| --- | --- | --- |
| **Data collection** |  |  |
| Space group | P2_1_2_1_2_1_ | P2_1_2_1_2 |
| Cell parameters  a, b, c, (*Å)* | 39.17, 69.24, 181.45 | 59.76, 105.21, 194.84 |
| Wavelength (*Å)* | 1.000 | 1.000 |
| Resolution limits (*Å)^a^* | 38.3 – 2.05 (2.17 – 2.05) | 39.0 – 1.85 (1.96 – 1.85) |
| Unique reflections | 31524 (4951) | 103844 (15707) |
| Completeness (%) | 98.7 (97.8) | 98.1 (93.1) |
| Redundancy | 3.60 (3.60) | 3.58 (3.26) |
| I/σI | 9.75 (1.81) | 11.16 (1.66) |
| R_merge_ (%) | 9.0 (72.5) | 8.0 (60.6) |
| CC (1/2) | 99.7 (80.3) | 99.9 (78.7) |
| **Refinement** |  |  |
| Resolution limits (*Å)* | 38.3 – 2.05 | 39.0 – 1.85 |
| R_cryst_ (%) | 23.6 | 20.1 |
| R_free_ (%) | 27.1 | 24.4 |
| Mean B value (*Å^2^)* | 46.1 | 30.7 |
| Ramachandran preferred/allowed/outlier regions (%) | 99.0/1.0/0.0 | 99.6/0.4/0.0 |
| **PDB accession code** | **9TRJ** | **9TRK** |

^a^ Values in parenthesis refer to the highest-resolution shell.
